# A protein aggregation-dependent scaling-free synaptic facilitation rule for neural network simulations

**DOI:** 10.1101/031856

**Authors:** Michele Sanguanini, Antonino Cattaneo

## Abstract

The regulation of mRNA translation at synaptic level is believed to be fundamental in memory and learning at cellular level. A family of RNA binding proteins (RBPs) which emerged to be important during development and in adult neurons is the one of Cytoplasmic Polyadenylation Element Binding proteins (CPEBs). *Drosophila* Orb2 (homolog of vertebrate CPEB2 protein and of the neural isoform of *Aplysia* CPEB) has been found to be involved in the translation of plasticity-dependent mRNAs and has been associated to Long Term Memory (LTM). Orb2 protein presents two main isoforms, Orb2A and Orb2B, which form an activity induced amyloid-like functional aggregate, which is thought to be the translation-inducing state of the RBP. Here we present a two-states continuous differential model for Orb2A-Orb2B aggregation and we propose it, more generally, as a new synaptic facilitation rule for learning processes involving protein aggregation-dependent plasticity (PADP).

## 1 Introduction

Different and complex cellular and molecular aspects underlie information learning and memory, even in simple invertebrate models such as *Aplysia californica* or *Drosophila melanogaster*^1^. A commonly used framework to model and schematize memory and learning phenomena is founded on the Hebb principle, which states that the synaptic connection between two neurons is facilitated if the neuronal pre-synaptic and post-synaptic activity correlates ^2^ consistently. A direct experimental validation of Hebbs principle is provided by the phenomenon of *Spike-time Dependent Plasticity*^3^ (STDP). From this principle, a commonly used STDP-based synaptic rule is derived for modeling and neuronal network simulations^4,5^, which increases or decreases the synaptic efficacy according to the time difference in the onset of the pre- and the post-synaptic activity. In general, synaptic learning rules should involve the activation of long-lasting mechanisms that are local in nature (to ensure local synaptic-specificity of synaptic plasticity).

A peculiar feature discovered in *Aplysia* neurons is the prion-like behavior of the neuronal isoform of the RNA binding protein CPEB^6,7,8,9^. The CPEB proteins are translational regulators of multiple mRNA species and they fall into two subfamilies. The CPEB1-like subfamily includes *mouse* CPEB1 protein and its orthologs *Xenopus* CPEB, *Drosophila* Orb protein and *Aplysia* CPEB^10,11^ (ApCPEB). These proteins bind the conserved U-rich Cytoplasmic Polyadenylation Element (CPE) sequence and mainly regulate the length of the poly-A tail–which is involved in the initiation of translation–of the target mRNA^10,11^. The CPEB2-like subfamily includes vertebrate CPEB2-4 proteins and the *Drosophila* Orb2 protein^10,11^. The proteins of this group don’t necessarily bind to a CPE sequence^10,12^ (there could be an important role of the secondary structure of the target mRNA^12,13^ in the recognition) and their mechanism of translation regulation is not directly linked to the polyadenylation state of the mRNA^10^. All CPEB proteins are present in two states, a translation inducing state and a repressor one, and the transition between these states is usually regulated (for example in mammalian CPEBs or in *Drosophila* Orb protein) through phosphorylation^14^. Some members of both CPEB1-like and CPEB2-like subfamilies, in particular ApCPEB^6,8,9^, *Drosophila* Orb2^15^ and the mouse isoform CPEB3^16,17^, show a peculiar transition process between the inducing state and the inhibiting one, which occurs through a conformational modification of the protein. The conformational modification occurs at the level of the Q-rich Intrinsically Disordered Domain (IDD) of the protein, which is located at the N-terminus and shows a prion-like behavior^6,8,9,15,16^–that is, once the CPEB domain undergoes the active state conformational change, it starts being a template for other CPEB proteins to change their IDD conformation, thus usually leading to protein aggregation.

ApCPEB presents an inactive-possibly translation-inhibiting-soluble state and an active, translation-promoting aggregated state^6^. These aggregates are self-sustaining, that is, once triggered, they grow and maintain themselves via autonomous biophysical properties, they are proteinase K resistant and positive to Thyoflavin T staining, characteristics that show their amyloid nature^6,8,9^. Studies in yeast have demonstrated that the aggregation occurs according to a protein-only-dependent mechanism, which is mediated by the Q-rich intrinsically disordered N-terminus of the protein^6,9^. A trigger for the aggregation of synaptic ApCPEB is the synaptic stimulation with serotonin ^7,8^, a neurotransmitter which has been found to be crucially involved in LTM in *Aplysia*^18^: indeed, the number of amyloid-like ApCPEB aggregates has been shown to correlate with memory retention tasks^8^.

*Drosophila* Orb2 protein, unlike the other fly CPEB protein Orb, shows a prion-prone N-terminus domain^8,10,19^, which triggers the formation of amyloid-like aggregates that are, again, self-sustainable, self-templating and functionally correlated with LTM^8^. The *orb2* gene undergoes alternative splicing and many Orb2 isoforms are produced ^10^-of these, two proteins of interest are the isoform A (Orb2A) and B (Orb2B)^8,10,19^. In particular, Orb2B is the most abundant isoform in the cell, is widely diffused and shows no or little aggregation propensity, while Orb2A is little expressed, is locally translated and aggregates quickly in proteinase- and SDS-resistant puncta^8^. The aggregation propensity of Orb2A makes this isoform a potential burden for the cell proteostasis and thus the Orb2A local level is strictly regulated by the cell through a finely-tuned translation-degradation equilibrium^20^. It has been shown that Orb2A mediates the local aggregation of Orb2B after synaptic stimulation, thus forming a heteromeric Orb2 aggregate^8,19^. It is reasonable to suppose that the heteromeric Orb2A-Orb2B aggregation follows a synaptic activity-dependent process of local Orb2A stabilization and accumulation ^20^. The two isoforms have complementary roles in LTM-related tasks. In fact, *Drosophila* mutants without Orb2B RNA-binding domain (RBD) show no LTM–when Orb2A RBD mutants show normal LTM^19^. On the contrary, Orb2A Q-rich N-domain is significantly involved in LTM, while Orb2B prion-like domain is not ^19^–yet a residual LTM is present in Orb2A IDD-deleted fly mutants, suggesting a minor role for activity-dependent Orb2B homomeric aggregation in Orb2 aggregate formation. It is likely that, after the IDD-mediated aggregation of Orb2(B), the protein makes a transition from a state of translation suppressor to a translation-permissive state-or even translation-facilitator, as suggested to occur for ApCPEB aggregates. The released mRNAs could then be involved in synaptic facilitation via multiple parallel processes: many mRNAs which are supposed to interact with Orb2 are involved in synaptic growth and functions^12^, so a single aggregation event could regulate the translation of the local transcriptome (and thus the local synaptic proteome and interactome) contemporarily at the level of many protein network nodes, providing a significant amplification step, while preserving long-lasting and locality properties.

We modeled Orb2 aggregation according to the available experimental data for Orb2A-Orb2B heteromeric aggregation^8,19^ (Figure 1A). At resting conditions, there are undetectable levels of Orb2A in the synapse^8^ and the Orb2A protein becomes detectable after synaptic stimulation. It has been shown that synaptic stimulation directly regulates Orb2A levels through the stabilizing effects 1) of the Orb2-interacting protein Tob and 2) of the phosphorylation mediated by the Tob-recruited kinase LimK and the delayed Orb2-destabilizing phosphatase activity of PP2A^20^, so it is reasonable to suppose that the degradation function depends on synaptic activity. The assumption that synaptic activity directly affects the translation rate of Orb2A is not trivial and there are no experimental data supporting it, so it won’t be considered. It has also been shown that the Orb2A isoform is necessary during acquisition and that the Orb2B isoform is required during memory consolidation^21^. Also, it seems that a neuromodulatory dopaminergic stimulation is necessary for both memory formation steps^21,22^.

**Figure 1:**
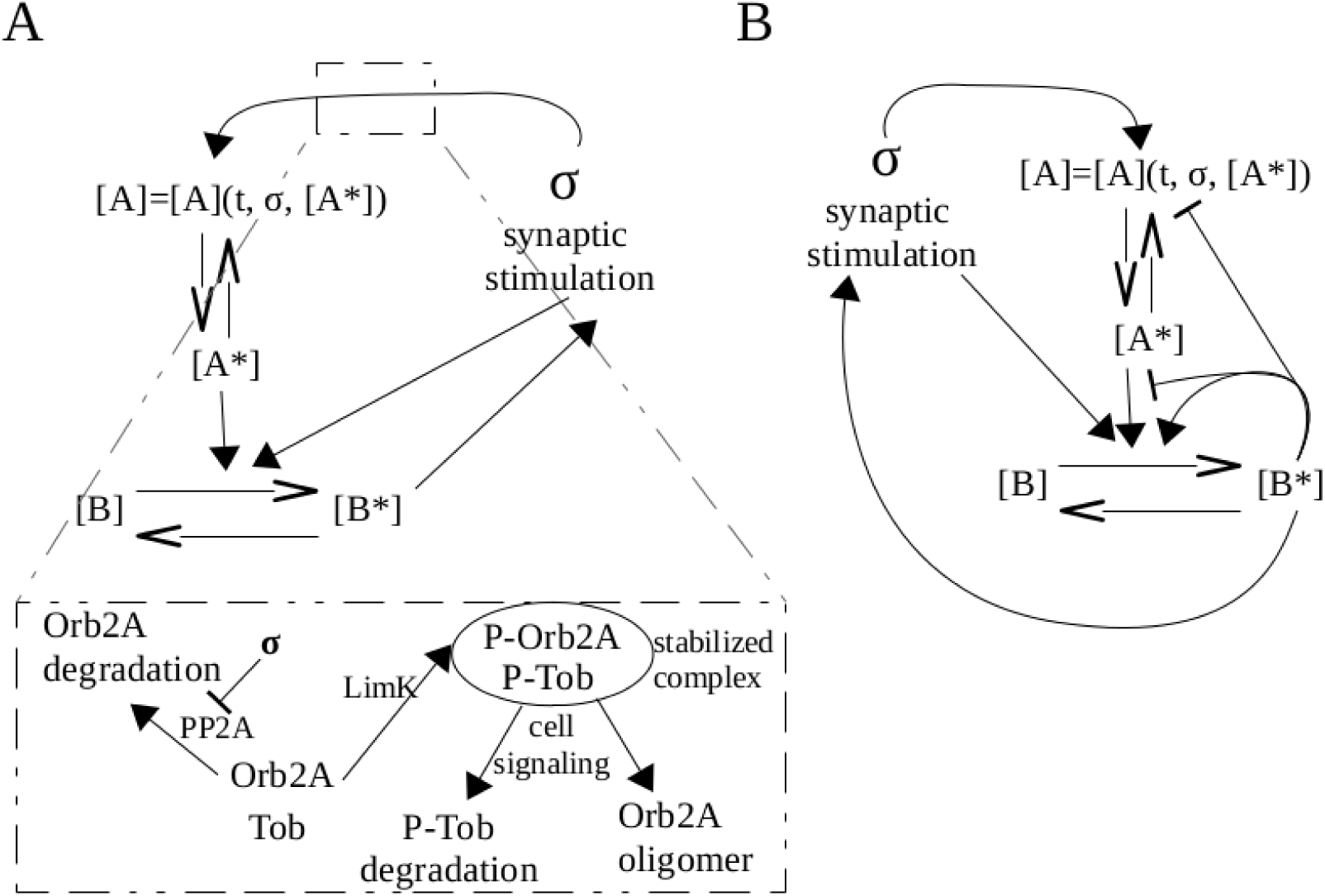
A) General scheme: the synaptic activity leads to an increase of Orb2A protein, which aggregates. The presence of Orb2A aggregate induces the hetero-oligomerization of Orb2B and thus the Orb2 aggregate increases the synaptic activity acting on the local synaptic translation. There is also a hypothesized residual Orb2B aggregation directly mediated by synaptic activity. The *embedded scheme* shows a possible mechanism of activity-dependent Orb2A aggregation *(adapted from* [^20^]). The synaptic activity leads to the suppression of PP2A phosphatase action, which destabilizes Orb2A, and the Tob-mediated recruitment of LimK. (Interestingly, unphosphorylated Tob protein is more stable than phospho-Tob.) It thus forms a stable phospho-Tob/phospho-Orb2A complex that will mature in the Orb2 aggregate, probably after the degradation of phospho-Tob and the conformational transition to amyloid-like form. B) Chemical reactions used to model the phenotype of the aggregation of *Drosophila* Orb2 protein.

The aim of this paper is to present a new synaptic facilitation rule, which is inspired by the physiological amyloid-like aggregation and involves the local protein translation at the synapse. An obvious advantage of this PADP mechanism is that it is self-regulating and contains a intrinsic saliency detector. After a significant synaptic stimulus which triggers the aggregation, the aggregate regulates its own growth and size, so the synapse does not incur in paradoxical aggregation and facilitation occurs with a shift of synaptic equilibrium to an increased steady-state which is not indefinitely high.

## 2 Self-sustaining protein aggregation general model

We now consider a general equation of local monomer production (the limiting species being in this case Orb2A) which includes the functions of local translation and local degradation:

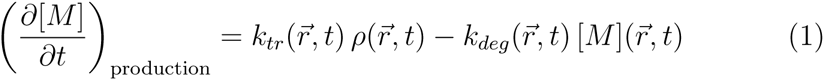

where the 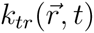 function describes the translation rate in space and time, 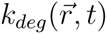 the degradation rate, 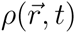 the local amount of Orb2A mRNA and 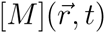 the local concentration of the monomeric protein.

Let’s now add a dependency of the monomer concentration on the synaptic activity. We assume that the monomer is produced locally after the synaptic stimulus, according to a function of the synaptic stimulus itself:

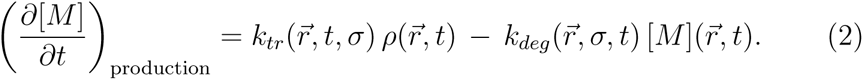

where *σ* is the synaptic stimulation time course.

The protein monomer is thus accumulated in a certain time window, which is governed by the parameters of the *σ*(*t*) function. The accumulation favors the transition to an aggregation-active state, that for Orb2 could be the formation of an Orb2A oligomeric nucleation seed and for ApCPEB could consist in the transition to the prion-like state. It is reasonable to assume that, in the cell, these molecular mechanisms are strictly regulated in a synaptic-activity dependent fashion: Orb2A is stabilized by interaction proteins like Tob and LimK^20^ and it’s possible that ApCPEB prion transition is locally regulated by synaptic activity-activated proteins (such as chaperones).

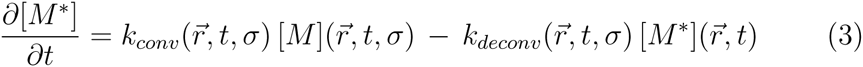

where the 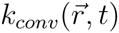 function describes the activation rate evolution, 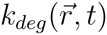 describes the rate of exit from the aggregation-prone state and 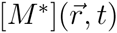 is defined as the concentration of monomer molecules that are in the aggregation-prone state.

When a certain amount of monomer protein enters the aggregation-prone state, it is likely that this seed catalyzes the prion-like conversion and thus the aggregation of more monomers:

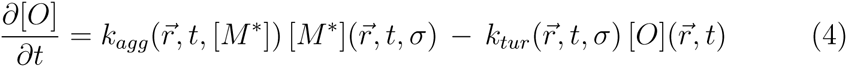

where [O] is the amount of monomers in aggregated state–regardless whether they form many short polymers, or few long ones, *k_agg_* is the rate of aggregation and *k_tur_* is the aggregate turnover rate. It is also plausible that the prion-like conversion rate is a function of the aggregation state *k_conv_* = *k_conv_* (…, [*O*]).

It has been shown that the size of ApCPEB aggregates in *Aplysia* cells is kept quite constant^8^, suggesting that there are mechanisms regulating locally the aggregation efficiency and providing a feedback mechanism limiting the size and number of aggregates to a maximum. It is reasonable to extend this reasoning to other functional aggregates. This would be due to aggregate-dependent changes in the disgregation functions so that *k_tur_* = *k_tur_*(*…*, [*O*]), and/or in the levels of free monomer itself ([*M*] = [*M*](…, [*O*])) – in the case of Orb2 aggregate. This is plausible because Orb2 RBP regulates also its own mRNA^12^.

It is now possible to write the general equations which describe the dynamics of a self-sustaining and self-limiting protein aggregate:

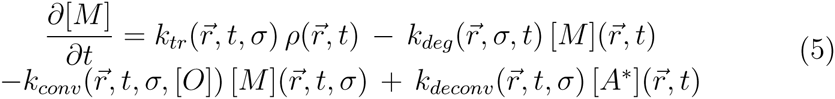

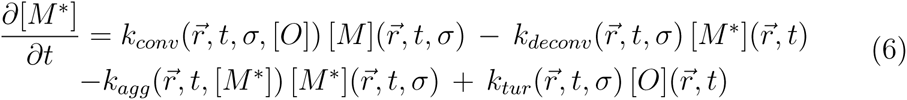

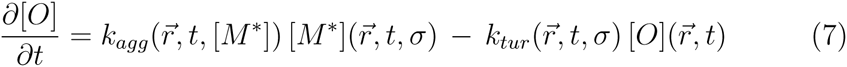

The aggregate [O] is the main output of the system, which affects the synaptic efficiency through the regulation of the local translation.

## 3 Protein-aggregation dependent synaptic rule (Orb2-inspired)

In order to model, in a first approximation, Orb2 aggregation we assume a series of conditions, starting from the previous general model. The first one is to consider the synapse as a punctiform compartment neglecting space variables and that there is no diffusion of the Orb2A protein outside the synapse. The second assumption is that the number of Orb2A mRNA molecules locally available for translation are constant; this is justified by growing evidence about a Orb2A mRNA post-transcriptional transition from a somatic non-protein coding to a local protein-coding form, which likely occurs at stimulated synapses thanks to the splicing mediator Nova^23^. From these conditions it comes that the studied variables are only functions of time and their variation is a total derivative according to *t*. The third hypothesis is that the aggregation seed is composed by Orb2A oligomers^19,20,21^, while the Orb2 mature aggregate is composed by the Orb2B isoform and, once formed, is completely self-sustaining. So, in our Orb2-inspired model, Equations 5 and 6 are not linked through the conservation of the monomer species. In partial accordance with Equation 6, we assume an (indirect) cross-dependence between the levels of oligomeric Orb2A and Orb2B aggregate. Again, we are considering the aggregate form to be a general state of the protein, without considering the number and/or the size of each individual aggregate. A fundamental assumption for the Orb2 model is that the Orb2A oligomer forms a seed for the further Orb2B aggregation^15,19,20,21^ and that–once Orb2 aggregate is grown to a certain level–the Orb2 oligomer becomes self-sustaining and invariant to the Orb2A seed presence. The Orb2A oligomer is thus possibly a synaptic tag to commit the activated synapse to long-term facilitation^21^. Interestingly, a monoaminergic secondary activity is needed for the further consolidation of memory^21,22^ (i.e. in the Orb2 system, the Orb2B isoform aggregation).

We assume that the local translation of Orb2A depends on the synaptic activity, according to the following equation:

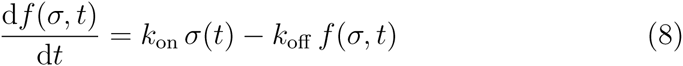

thus, the system response to the synaptic activity (in this case modeled as a binary function *σ*(*t*), which could be associated with the ON state of a minimum amount of the local monoamine channel population) grows with a *k*_on_ constant and becomes saturated and unresponsive with a *k*_off_ rate. The local levels of Orb2A monomer increase after synaptic activity^15,20^ and this growth is regulated by the stabilizing effect of Tob/LimK-mediated phosphorylation of Orb2A and by the destabilization of phosphorylated Orb2A interacting protein Tob^20^: the superposition of these processes likely identifies a time window for synaptic accumulation and oligomerization of Orb2A. Also, if Orb2A is to be considered a synaptic tag, it’s likely that the Orb2A production should become insensitive to further synaptic activity after the long-term consolidation of memory–or the formation of Orb2B amyloid-like oligomer which potentially ensures an indefinitely long-lived synaptic facilitation.

The Equation 9 tries to include the described properties of Orb2A translation:

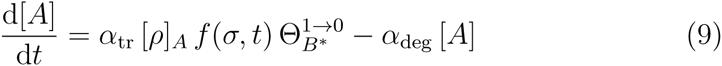

where *α*_tr_ is the local translation rate of Orb2A, [*ρ*]*_A_* is the concentration of Orb2A mRNA–assumed to be constant, *α*_deg_ is the local degradation rate and 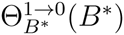 is a Heaviside-like continuous function which assumes values close to zero when the amount of Orb2 aggregate approaches a threshold value [*B*]*_θ,A_* and close to one for smaller values.

Orb2A has a strong intrinsic tendency to aggregation^15^, so we ignore the transition to an aggregation prone state of the monomer (Equation 3) and directly model the Orb2A aggregation:

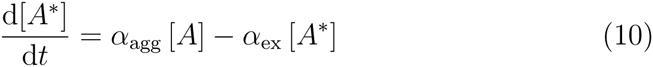

where *α*_agg_ is the aggregation rate and *α*_ex_ is the aggregate exit rate. Applying the conservation of Orb2A species to Equations 9 and 10, we obtain:

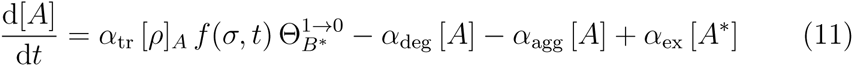

The Orb2A/Orb2B hetero-oligomerization, which initiates the formation of Orb2 aggregates that correlate with long-term memory, depends on both the presence of an Orb2A aggregation seed^15,19^ (i.e. oligomer) and of synaptic stimulation^21^. Given that the translation-dependent L-LTM is not induced by an intensive (i.e. non-spaced) training^24^ and that *Drosophila* Orb2-dependent LTM easily occurs after a series of stimulations and recoveries ^10^, a good candidate for this synaptic activity is the hour-long rhythmic dopaminergic stimulation seen by Plagais and colleagues after spaced learning ^25^, which has been shown to inhibit anesthesia-resistant memory (ARM) and thus to gate LTM^25^. It has been shown–at least for *Aplysia* CPEB amyloid-like aggregates–that the dimensions of the higher-order aggregates (also called *puncta*) are kept at a steady level^8^, with a turnover rate of about 20% in 48h. This steady-state behavior could be applied also to Orb2. Also, Orb2 aggregate self-sustaining steady state appears to be independent of the presence of Orb2A–after reaching the mature state. In order to model this complex behavior, we consider a first simplification: both the mRNA and the protein of Orb2B are present at high levels in the cytoplasm, so we assume the translated non-aggregated Orb2B isoform is kept constant at the dendrite level, because the local levels of the protein are instantaneously buffered against a much larger somatic reservoir.

The proposed equation for Orb2B aggregation is the following:

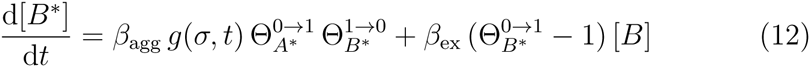

where [B]* is the amount of Orb2B in aggregated state, *β*_agg_ is the Orb2A seed-dependent aggregation rate and includes the (constant) Orb2B monomer level, *β*_ex_ rules the aggregate dissociation when the oligomer doesn’t reach the steady level and g(*σ*) is the synapse reaction to a (binary) synaptic stimulation. 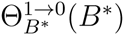 is a Heaviside-like continuous function which assumes values close to zero when the amount of Orb2 aggregate approaches a threshold value [B]*_θ,BA_* and close to one for smaller values: that is, the growth of Orb2B aggregate becomes independent to Orb2A seed after reaching a threshold value. An analogous behavior, but opposite, is true for 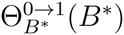 (self-sustaining steady state at [*B\**]*_θ_,_BB_*) and 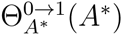 (responsiveness to the Orb2A oligomer [*A\**]*_θ_,_min_*). The Equation 12 doesn’t take into account the possible weak homoinduced Orb2B aggregation, which has been hypothesized in order to explain the residual LTM observed in Orb2A-only defective flies^19^. Since it’s not been established which are the stimulation dependent parameters of functions *f*(*σ*) and *g*(*σ*), we are assuming that *f*(*σ*) = *g*(*σ*) and that Orb2A and Orb2B are both responsive to the same dopaminergic stimulus *σ*(*t*).

## 4 Facilitation rule

The PADP synaptic rule states that the synaptic efficiency scales with the amount of Orb2B aggregate; as we modeled the aggregation, after reaching an activity-dependent threshold the aggregate self-templates itself. These conditions imply an all-or-nothing response of the synapse: if a salient condition has been presented for enough time to trigger the formation of an amyloid aggregate, after a reasonable time interval between the aggregation and the consequent translational activation of pro-plasticity factors (called here *latency time τ*), the synapse will make a transition to a higher efficiency stable state. We modeled the PADP synaptic efficiency update with a sigmoidal function (**Figure 2**) because it has a steep upper-bounded behavior that is well-suited for the postulated almost two-state plasticity rule,

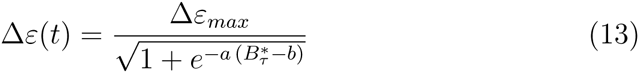

where Δ*ε_max_* is the maximal ΔEPSP (Excitatory Post-synaptic Potential) found at the synapse after an experimental facilitation protocol, *a, b* and *c* are the parameters that regulate the sigmoid function behavior, 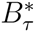 is the composed function [*B\**] o *T* where

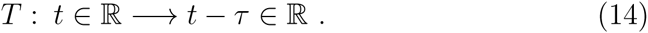

**Figure 2:**
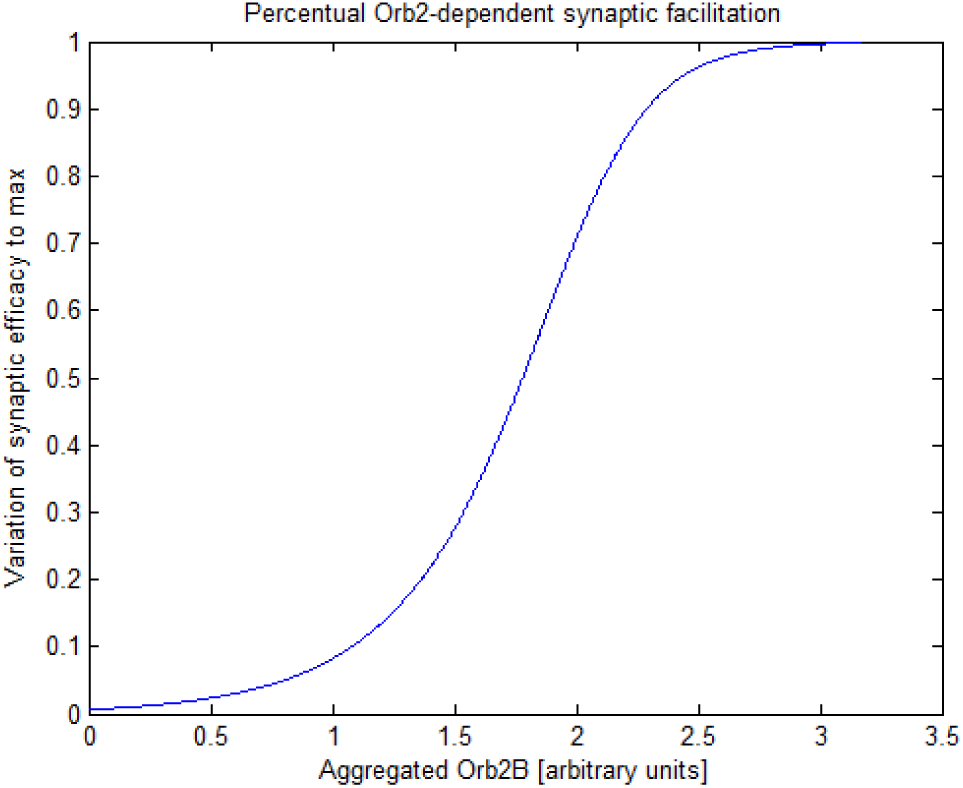
Plot of synaptic efficiency variation (% ratio to Δ*ε_max_*) according to aggregate Orb2B concentration (arbitrary units). a=5, b=2.

In this way, also the latency time between the aggregation trigger and the facilitation onset is taken into account.

## 5 Numerical simulations

Numerical simulations of Equations 10, 12 and 15 were made using a Runge-Kutta fourth-order algorithm with an integration time step of 0.001 seconds and the parameters shown in **Table 1**. The Heaviside-like functions which were used for the simulations are continuous stepped functions:

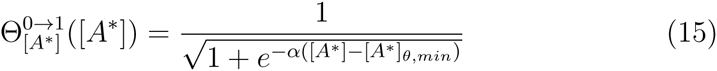

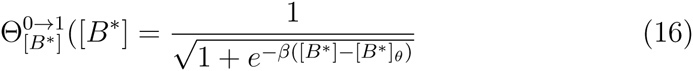

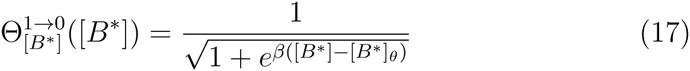

where *α, β ∈* ℕ: *α, β* ≫ 1 in order to get a steep transition at the threshold and [*B\**]*_θ_* = [*B\**]*_θ_,_A_* = [*B\**]*_θ_,_BA_* = [*B\**]*_θ_*,*_BB_* because we assume only one level of Orb2 aggregate-dependent regulation of Orb2A and Orb2B aggregation.

**Table 1:** Parameters of Orb2 model used in numerical simulations.

| Numerical values of parameters |  |  |  |  |  |
| --- | --- | --- | --- | --- | --- |
| $k_{\text{on}}$ | 1 | Hz | $k_{\text{off}}$ | 0.7 | Hz |
| $\alpha_{\text{tr}} \cdot [\rho]_A$ | 0.005 | $\frac{\text{units}}{s}$ | $\alpha_{\text{deg}}$ | 0.002 | Hz |
| $\alpha_{\text{agg}}$ | 0.008 | Hz | $\alpha_{\text{ex}}$ | 0.001 | Hz |
| $[A^*]_{\theta}$ | 3 | units | | | |
| $\beta_{\text{agg}}$ | 1 | $\frac{\text{units}}{s}$ | $\beta_{\text{ex}}$ | 0.0004 | Hz |
| $[B^*]_{\theta}$ | 3 | units | | | |
| $\alpha$ | 50 | | $\beta$ | 50 | |

The stimulation pattern *σ*(*t*) was a unit square wave function with 30% duty cycle and period T=5s. (The *duty cycle* of a square stimulus is the ratio of the period which is occupied by the stimulation.)

We then tested the model in order to assess: 1) the sensitivity to the length of the stimulation–that is, a short, spurious stimulation shouldn’t trigger the long term facilitation–and 2) the self-limited growth even after a continuous stimulation. Figure 3 shows a simulation with 4000 s of stimulation time (a physiological-like contest) and the self-limiting behavior is visible. Figure 4 shows a simulation with 2000 s of stimulation time: since Orb2A levels haven’t reached a sufficient level to form the aggregation seed, no aggregated Orb2B forms. Figure 5 shows a simulation with continuous stimulation: the aggregated Orb2B level reaches a steady-state dimension similar to the physiological one.

**Figure 3:**
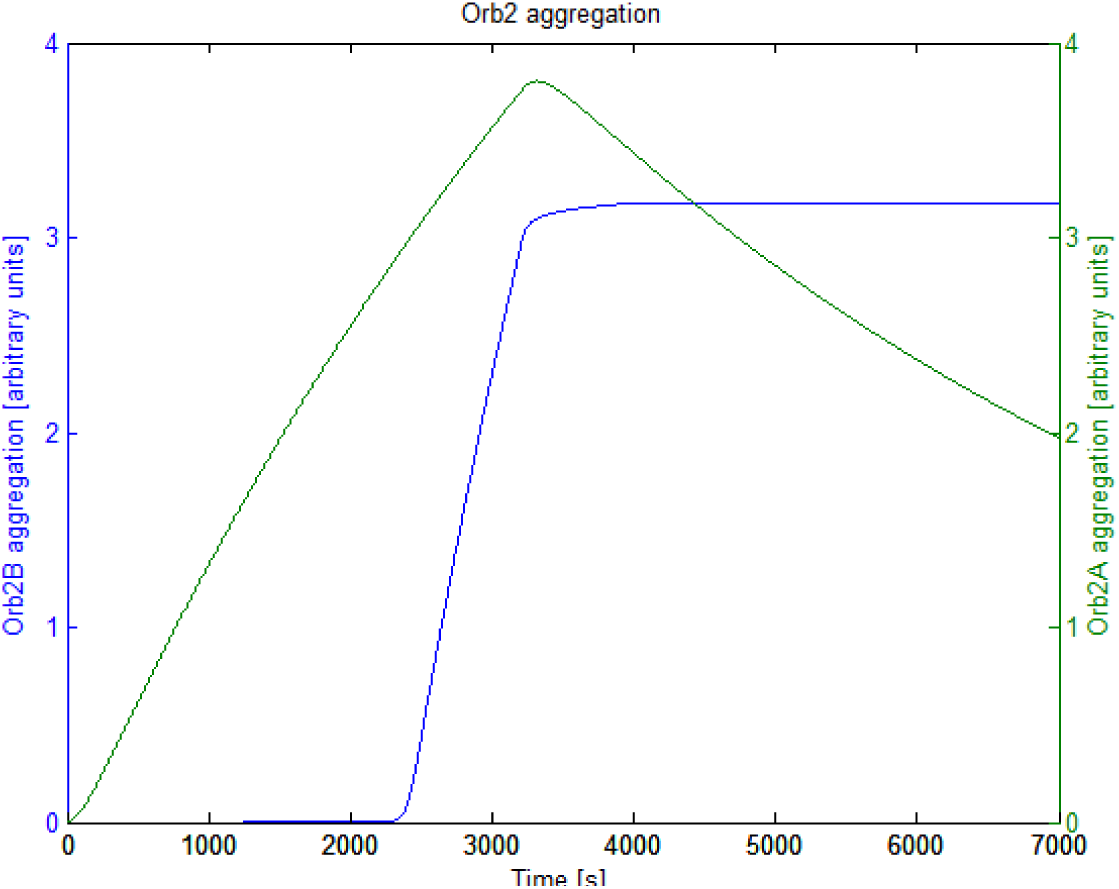
Simulation of the aggregation levels of Orb2B (blue) and Orb2A (green) with 4000s of binary square stimulation (T=5s, duty cycle 30%).

**Figure 4:**
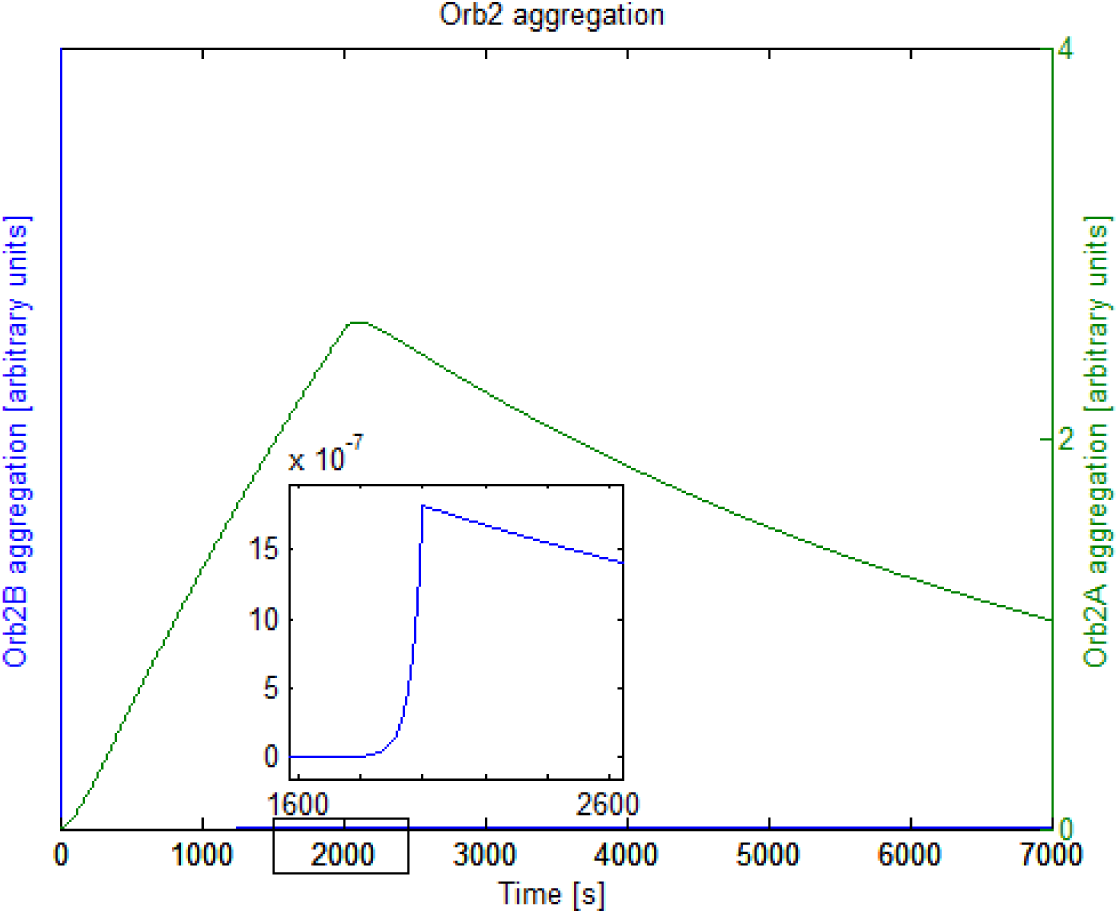
Simulation of the aggregation levels of Orb2B (blue) and Orb2A (green) with 2000s of binary square stimulation (T=5s, duty cycle 30%).

**Figure 5:**
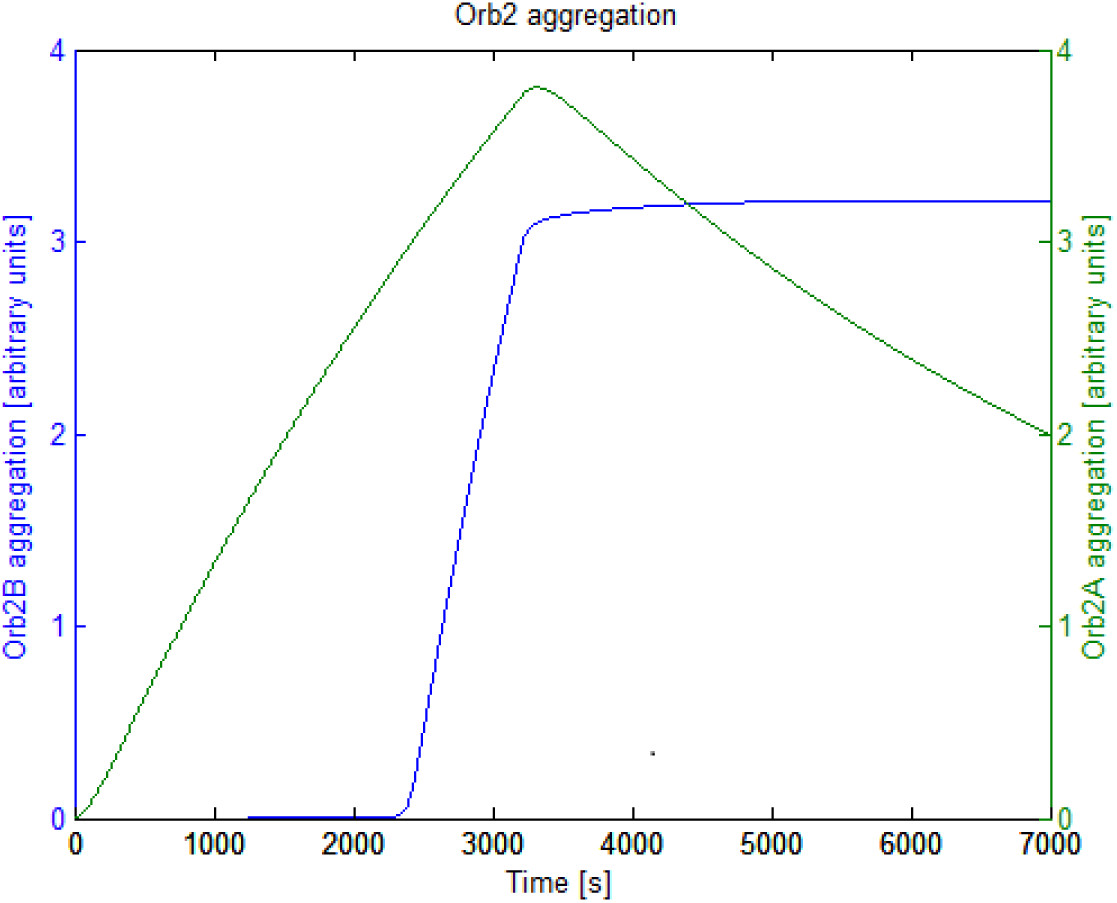
Simulation of the aggregation levels of Orb2B (blue) and Orb2A (green) with continuous binary square stimulation (T=5s, duty cycle 30%).

## 6 Dependency on the characteristics of the stimulus

We then asked how the model responds to different characteristics of the (binary) stimulus *σ*(*t*). The numerical simulation were made using a 4th order Dormand-Prince solver with variable time step between 0.001 and 0.01 seconds; the stimulation functions were forced to zero after 4000 seconds (to assume a probably physiological-like environment). We first considered the dependency on the presentation “frequency” of a constant length square stimulus (in Figure 6, 1.5 seconds), which has been produced using a square function with varying period and duty cycle, in order to get a 1.5s stimulation followed by a uniformly distributed resting time. From Figure 6 it can be seen that there is a threshold of frequency (in our model being between 0.14 and 0.2 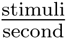) which leads to the self-sustaining aggregation. It could be seen that the simulated stimulus has two different characteristics: the *period* of stimulus presentation (T) and the *duty cycle*, which is more generally a parameter of the time when the system is able to produce a maximal ON signal compared to the relaxing time (here it will be called *mean effective stimulation time*).

**Figure 6:**
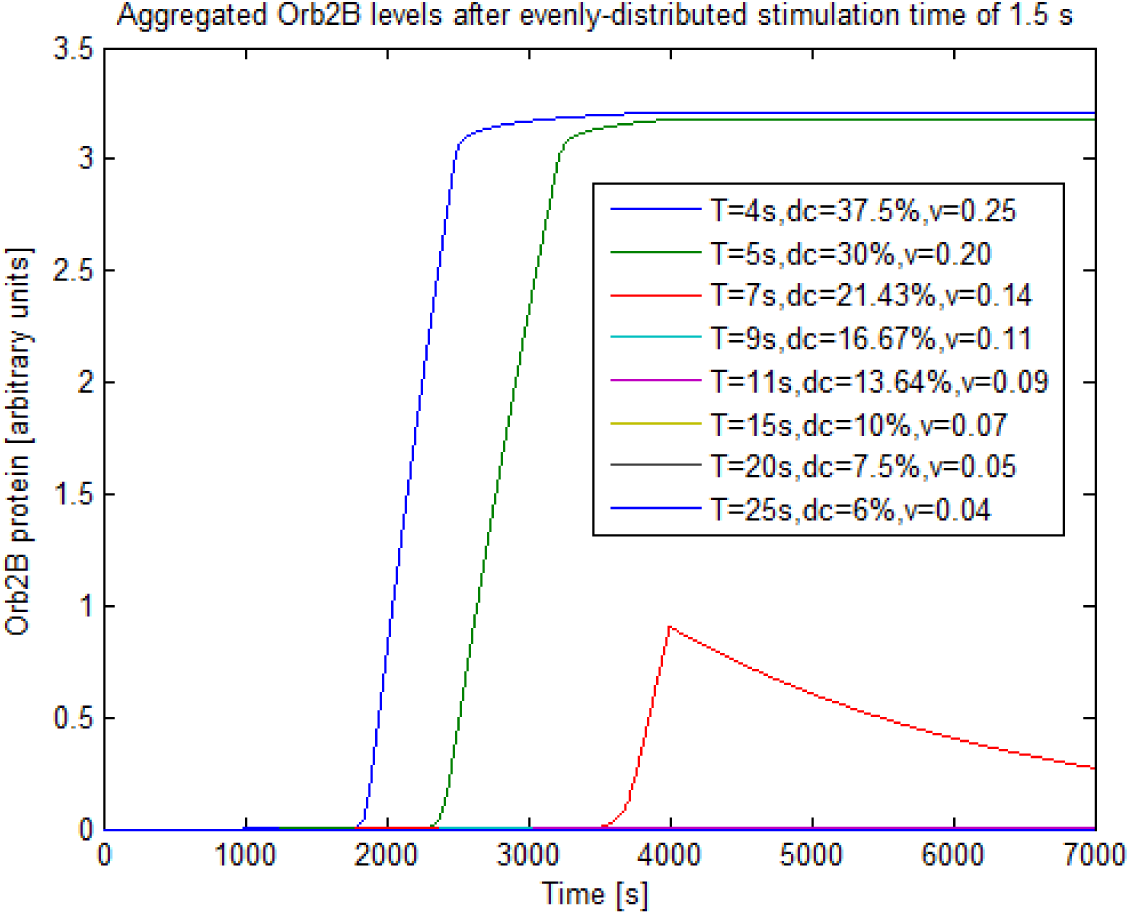
Simulation of the levels of aggregated Orb2B with fixed stimulation time of 1.5s and uniformly distributed variable resting time.

Figure 7 A shows a multiple simulation where the period was the same (5 seconds) and the duty cycle was changed and Figure 7 B shows a multiple simulation where the duty cycle was kept constant (27%) and the period was variable–a continuous stimulation and a random one (Bernoulli binary distribution, p=0.27) were used as a control. The simulations show that the system, as we modeled it, is sensitive to the mean effective stimulation time rather than the period of stimulation and that the responsiveness of the aggregation to a given duty cycle depends on the total time of stimulation. That is, if we consider again Figure 6, a rhythmic stimulation with frequency of 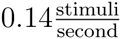 which ceases after 4000s doesn’t trigger the self-sustaining aggregation of Orb2B, but a prolonged stimulation could trigger the oligomerization (Supplementary figure 1).

**Figure 7:**
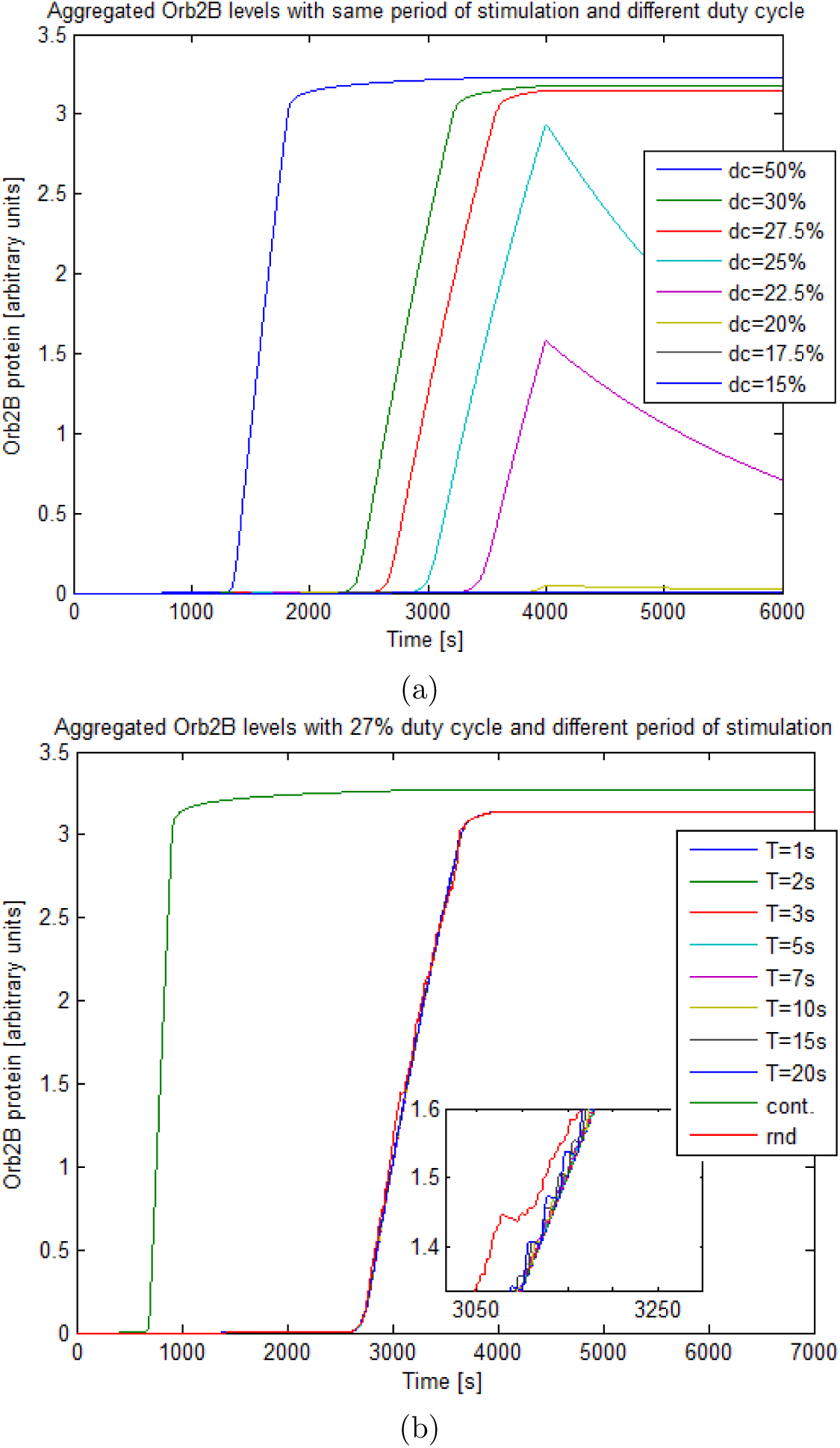
Multiple simulations of the levels of aggregated Orb2B with a) constant period T of 5s and variable duty cycle and b) constant duty cycle of 27% (close to the stabilization threshold) and variable period. There is a neat dependency of the aggregation of Orb2B from the mean effective stimulation time.

The shown Orb2-inspired aggregation system reveals an intrinsic ability to sustain a long-term summation of the external stimuli and could act as an integrator of the past activity of the synapse. The summation could occur at the level of the stimulus function *f*(*σ*), of the Orb2A-Orb2B oligomerization or at both levels. As it could be expected, Supplementary figure 2 shows that the function *f*(*σ*) sums the binary stimulus in continuous patterns. In order to assess the intrinsic summation ability of Orb2A-Orb2B we set *f*(*σ*) = *σ*(*t*) and then tested the sensitivity of the system to both period (Supplementary figure 3) and duty cycle (Supplementary figure 4) of the pure binary square stimulation. The simulations show that the modeled aggregation system can act as an intrinsic integrator of the synaptic stimuli.

## 7 Discussion

We presented a model of continuous differential equations for the Orb2-mediated mechanisms of synaptic plasticity. This synaptic plasticity mechanism has been shown to be important for a paradigm of LTM in Drosophila, but the interest of aggregation-dependent synaptic plasticity goes beyond this specific example and could represent a more general mechanism. The simulations performed show that the model is consistent with the molecular behavior of Orb2 aggregates seen *in vitro* and *in vivo*. Orb2 protein aggregation is likely to be a detector of salient (i.e. biologically important) activity and a self-sustaining and self-limiting memory device whose outcome is very reliable and stable.

However, it is necessary to point out some limitations arising from such a model, because of the simplifying assumption we had to choose. Being a continuous model and considering the two states of the protein (free and aggregated) as two “compartments”, the model does not take into account the probabilistic aspects of the local aggregation (for example the sensitivity to the fluctuations of protein concentration). Fluctuations in protein concentration might be particularly relevant, for the small volume and low protein numbers in the volume of a dendritic spine. Also, the model does not describe other kinetic aspects such as aggregate polarisation, the general mechanism of aggregation^26^ and so on. Given the little knowledge of molecular interacting partners of the aggregate and the pathways of cellular regulation of Orb2 aggregates ^20,27^ and the lack of a rigorous kinetic study of Orb2A-Orb2B homo- and hetero-induced aggregation, we found to be more conservative to model the overall known phenotype of Orb2 aggregate rather than to guess the molecular-level behaviour of the involved proteins. Other aspects of the model-now set as constant or null for simplicity sake–that will have to be considered in future implementations of the model are the transition from a point synapse to a spatially extended one and a more complex dynamic of Orb2A and Orb2B mRNAs and proteins, including diffusive components, RNA degradation, local depletion upon translation or aggregation, and so on. It would be surely significant to investigate a differential diffusivity between Orb2A or Orb2B free proteins and Orb2 aggregate–which *in vivo* probably characterizes the local (synaptic) specificity of PADP.

Another focal point in our model is the behavior of the aggregation according to the stimulation pattern. Using a binary periodic stimulation, we have shown that our system detects the likelihood of stimulation in a given time window, rather than the frequency of the stimulation *per se*. It has been shown that, during a spaced LTM-related training, there is the rise of a hours-long rhythmic dopaminergic stimulation which has a more uniform frequency spectrum and a higher amplitude than the basal state and could gate the transition between ARM and LTM^25^. If our assumptions are correct, the model would predict that the rhythmic stimulation should be permissive for the transition to LTM and the frequency of the stimulation wouldn’t be instructive in this process, but it would be involved in reaching a sufficient level of stimulation during Orb2 aggregation time window. From an experimental point of view, this hypothesis could be assessed through, for example, the optogenetic control of specific populations of dopaminergic neurons in the mushroom body. Also, our model predicts that the Orb2 physiological aggregates at the synaptic level function as a high-pass filter which would discriminate the stronger, experience-evoked rhythmic stimulation from the spurious ones.

A further characteristic which emerged from the modeling is that the seeded aggregation process has an intrinsic ability to integrate the synaptic activity through time and locally. If this property was experimentally confirmed, the Orb2A-Orb2B system would extend the ability of synapses to integrate the information from the classic spatial cooperativity to a more complex spatio-temporal pattern.

## 8 Conclusions

The evidence of the Orb2 protein-only mechanisms involved in LTM could be used as an alternative to common synaptic facilitation rules such as the STDP^5^ or the *Aplysia*-inspired activity-dependent presynaptic facilitation–ADPF^28^. STDP and ADPF are not self-regulating mechanisms, so they need extra-synaptic scaling mechanisms in order not to reach the synaptic saturation. The PADP does not incur in these limitations because it is based on a self-limiting aggregation process and lends itself to be heuristically applied in biological- and bioengineering-derived neural network simulations.

To our knowledge, this model is the first one which takes into account both protein aggregation as well as the local translation during synaptic facilitation processes. This approach could be further applied to other synaptic plasticity phenomena which include local translation, such as the BDNF-induced plasticity^29^ or mTORC-dependent plasticity^30^. There are other phenomena which are mediated by the formation of a stably active biological entity-just like the Orb2 functional amyloid–and are involved in memory induction and/or maintenance. A paradigmatic case, albeit controversial, is the kinase PKMζ, which is thought to be involved in memory long-term maintenance ^31^. PKMζ is locally translated after a salient synaptic activity and is constitutively active (it doesn’t rely on a secondary messenger to maintain its activity), in a way which is similar to the Orb2 aggregates here presented. Equations 5, 6 and 7 could be easily used as a starting point for a generalization of PADP in a plasticity rule which depends on synaptic protein network modification through local translation.

## Acknowledgements

We would like to thank prof A. Bazzani for useful discussions.

